# SBT-272 improves TDP-43 pathology in the ALS motor cortex by modulating mitochondrial integrity, motility, and function

**DOI:** 10.1101/2022.10.04.510854

**Authors:** Mukesh Gautam, Barış Genç, Benjamin Helmold, Angela Ahrens, Janis Kuka, Marina Makrecka-Kuka, Aksu Günay, Nuran Koçak, Izaak R. Aguilar-Wickings, Dennis Keefe, Guozhu Zheng, Suchitra Swaminathan, Martin Redmon, Hatim A. Zariwala, P. Hande Özdinler

**Author notes:** To whom correspondence should be addressed: P. Hande Ozdinler.

## Abstract

Mitochondrial defects are one of the common underlying causes of neuronal vulnerability in motor neuron diseases, such as amyotrophic lateral sclerosis (ALS), and TDP-43 pathology is the most common proteinopathy in ALS. Disrupted inner mitochondrial membrane (IMM) reported in the upper motor neurons (UMNs) of ALS patients with TDP-43 pathology is recapitulated in the UMNs of well-characterized mutant hTDP-43 mouse models of ALS. The construct validity, such as common cellular pathology in mice and human, offers a unique opportunity to test treatment strategies that may translate. SBT-272 is a well-tolerated brain-penetrant small molecule that stabilizes cardiolipin, a phospholipid found in IMM, thereby restoring mitochondrial structure and respiratory function. We investigated whether SBT-272 can improve IMM structure and health in UMNs diseased with TDP-43 pathology in our well-characterized UMN reporter line for ALS. We found that SBT-272 significantly improved mitochondrial structural integrity and restored mitochondrial motility and function. This led to improved health of diseased UMNs *in vitro.* In comparison to edaravone and AMX0035, SBT-272 appeared more effective in restoring health of diseased UMNs. Chronic treatment of SBT-272 for sixty days starting at an early symptomatic stage of the disease *in vivo* led to a reduction in astrogliosis, microgliosis, and retention of UMN degeneration in the ALS motor cortex. Our results underscore the therapeutic potential of SBT-272, especially within the context of TDP-43 pathology and mitochondrial dysfunction.

**Highlights:**

1. Early and progressive upper motor neuron (UMN) degeneration defines ALS pathology
2. Mitochondrial defects are prominent and common in UMNs with TDP-43 pathology
3. SBT-272 treatment improves mitochondrial stability, mobility and function
4. SBT-272 treatment reduces astrogliosis, microgliosis and improves UMN health

## Introduction

Upper motor neurons (UMNs) are one of the key components of motor neuron circuitry, as they have a unique ability to collect synaptic input from many different types of cortical neurons and to relay the integrated information to spinal cord targets with very high specificity (Eisen, 2021; Lemon, 2008; Lemon, 2021). Their degeneration is a hallmark of many motor neuron diseases, including amyotrophic lateral sclerosis (ALS) (Brown and Robberecht, 2001; Brunet et al., 2020; Charcot and Marie, 1885; Fink, 2013; Gunes et al., 2020). Despite their importance and clinical relevance, UMNs have not generally been addressed sufficiently in pre-clinical studies of ALS and other motor neuron diseases. However, there is now ample evidence showing the relevance of improving UMN health as a potential treatment strategy. UMN degeneration occurs early in the disease (Geevasinga et al., 2015; Geevasinga et al., 2016; Thomsen et al., 2014), and is not necessarily be a function of spinal motor neuron degeneration (Eisen, 2021; Genc et al., 2022b; Marques et al., 2021). The generation and characterization of a novel UMN reporter line, in which UMNs can be distinguished among other cortical cells by an eGFP expression that is stable and long-lasting (Yasvoina et al., 2013), now enables cell-type specific analysis of UMNs with cellular precision that was not possible before.

TDP-43 pathology stands out as one of the most commonly observed proteinopathies in ALS patients (Arai et al., 2006; Neumann et al., 2006). Mutations in TARDP gene, which codes for the TDP-43 protein, have been linked to ALS (Buratti, 2015; Buratti and Baralle, 2008; Cykowski et al., 2017; Mackenzie et al., 2007; Neumann et al., 2006). Notably, ALS patients who do not have mutations in the TARDP gene also display protein aggregates that include a phosphorylated form of TDP-43 in the cytoplasm and/or nucleus of neurons (Ling et al., 2013), suggesting that the pathology widely occurs regardless of mutation. Therefore, understanding and resolving TDP-43 pathology will have broad implications and has been an important quest.

TDP-43 is a DNA/RNA-binding nucleoprotein that plays a role in various cellular functions. It shuttles back and forth between the cytoplasm and nucleus and steady-state levels are maintained by self-regulatory mechanisms (Ayala et al., 2008). TDP-43 pathology is observed in the central nervous system of non-ALS patients as well, including ALS with frontotemporal lobar dementia (ALS-FTD), frontotemporal lobar degeneration with TDP-43 inclusions, and Alzheimer’s disease (Cotelli et al., 2013; Ling et al., 2013; Wilson et al., 2011).

One of the features of UMNs diseased with TDP-43 pathology in both sporadic and familial ALS patients is at the site of the mitochondrion. The mitochondria of UMNs in these patients have disrupted inner mitochondrial membrane (IMM), while the outer membrane remains intact. This IMM defect is localized to the UMNs, as other neurons do not exhibit this mitochondrial defect (Gautam et al., 2019a).

Even though they are in different species, the UMNs of prp-hTDP-43^A315T^ mice, – developed to mimic human TDP-43 pathology–, and UMNs of ALS patients with TDP-43 pathology display very similar mitochondrial problems along the IMM (Gautam et al., 2019a; Wang et al., 2019; Wang et al., 2016). These findings have two implications: first is the realization that translation is at the cellular level, and second is the importance of maintaining IMM stability and integrity as a potential treatment strategy to improve the health of UMNs, especially in ALS patients with TDP-43 pathology.

Maintaining the integrity of IMM is important for nearly every aspect of mitochondrial function, including establishing the voltage gradient required for ATP generation (Kuhlbrandt, 2015; Paumard et al., 2002), for influencing mitochondrial motility/dynamics (Lin and Sheng, 2015; Miller and Sheetz, 2004; Verburg and Hollenbeck, 2008), and for the exchange of metabolites, ions, and nucleotides across the IMM. Cytoplasmic leakage of mitochondrial components, including mtDNA, is known to initiate neuroimmune reactions (Riley and Tait, 2020; West and Shadel, 2017) and contributes to TDP-43 pathology in the motor cortex. Therefore, a compound that can restore IMM integrity in UMNs may significantly contribute to the overall health and function of these diseased neurons. One such approach is stabilizing the IMM cardiolipin, a phospholipid that influences the IMM morphology and the assembly of IMM proteins (Chicco and Sparagna, 2007). The presence of abnormal cardiolipin, i.e., either structurally modified or aberrantly localized, has been associated with impaired neurogenesis, aging, and neurodegenerative diseases (Falabella et al., 2021). Cellular stress due to disease causing mutations or protein aggregates can remodel cardiolipin via stress-induced lipid peroxidation or impact its metabolism (Camilleri et al., 2020; David et al., 2005). Therefore, cardiolipin might also be crucial for establishing the integrity and stability of IMM in the mitochondria with respect to TDP-43 pathologies. However, cardiolipin modification has not been systematically evaluated in neurons with mutant TDP-43 cells, albeit widespread mitochondrial dysfunction is established (Davis et al., 2018; Gao et al., 2019).

Recently, drug discovery studies identified a novel cardiolipin targeted therapeutic molecule, SBT-272. A closely related compound, elamipretide binds to and stabilizes cardiolipin (Mitchell et al., 2020). Therefore, we investigated whether SBT-272 treatment would restore IMM integrity in diseased UMNs. We tested whether that would translate into mitochondrial functional improvements using quantitative methods, selectively within the context of the brain component of ALS with TDP-43 pathology. Our findings reveal a therapeutic effect of SBT-272 treatment for ALS, and perhaps also for diseases in which mitochondrial problems and IMM disruption are major contributing causes of neuronal vulnerability and degeneration.

## Materials and Methods

### Animals

#### Mice

All animal procedures were approved by the Northwestern University Animal Care and Use Committee and comply with the standards of the National Institutes of Health. All mice were in C57BL/6J background. UCHL1-eGFP mice were generated in the Ozdinler Laboratory (Yasvoina et al., 2013) and are now available at the Jackson Laboratory (#022476). Hemizygous UCHL1-eGFP females mated with hemizygous prp-hTDP-43^A315T^ males (Wegorzewska et al., 2009) (Jackson Laboratory, #010700) to generate prp-hTDP-43^A315T^-UeGFP mice. prp-hTDP-43^A315T^ mice were fed with gel diet (DietGel76A, ClearH_2_O, ME), to eliminate gastrointestinal problems. Transgenic mice were identified as described (Gautam et al., 2019a; Wegorzewska et al., 2009).

#### Rats

All studies were performed in male hsd:Sprague Dawley^®^ rats procured from Envigo at 275– 300g weight. All animal procedures were approved by LIOS Animal Care and Use Committee.

### Identification and characterization of SBT-272

SBT-272 was identified and characterized by Stealth BioTherapeutics (Needham, MA). SBT-272 was dissolved in saline at a concentration of 10mg/ml. For rodent pharmacokinetic studies, the animals were dosed at 1ml/kg volume.

#### SBT-272 in anoxic permeabilized rat cardiac fibers

Permeabilized cardiac fibers were prepared from the left ventricle of the normoxic heart as reported (Kuka et al., 2012). To induce anoxia, the maximal respiration rate of a sample was stimulated by the addition of succinate 10mM plus rotenone 0.5μM and ADP 5mM, and the preparation was left to consume all the oxygen in a respiratory chamber (10–20 minutes in O_2_k respirometers, Oroboros^®^ Instruments) (Makrecka et al., 2014). After 30 minutes of anoxia, oxygen was reintroduced to the chamber. Once the oxygen concentration equilibrated to initial levels, the chamber was closed and oxygen flux was monitored for 10 minutes. H_2_O_2_ flux was measured simultaneously with respirometry in the O_2_k fluorometer using the H_2_O_2_-sensitive probe Ampliflu™ Red (Makrecka-Kuka et al., 2015). SBT-72 or vehicle was added before the introduction of permeabilized fibers to the chamber.

#### Rat model of acute kidney injury induced by ischemia-reperfusion

Acute kidney injury was induced by clamping the renal artery for 30 minutes before allowing reperfusion of blood in isoflurane-anesthetized male rats (*n*=8/group). Blood (for plasma) was drawn through the jugular vein twice (before ischemia induction and 24 hours after ischemia reperfusion). Creatinine and blood urea nitrogen were measured in plasma. Test compounds were administered 30 minutes before ischemia (2mg/kg SC, 2mg/ml prepared in saline) and again 5 minutes before reperfusion (2mg/kg SC).

#### Brain mitochondrial respiration in rats

Male rats were treated with two doses of SBT-272 (1 and 5mg/kg SC) once at 48 hours and 28 hours prior to decapitation. The rats were also treated with ET-1 or vehicle through stereotactic surgical route 24 hours before decapitation. The brains were isolated after decapitation. Stroke or sham-injured brain region was sliced (2.0mm thick) for mitochondrial function evaluation using high-resolution respirometry (Oroboros Instruments). Tissues were weighed and Mir05 buffer supplemented with 20mM creatine was added at a concentration of 80mg/ml (wet weight per ml). Tissues were homogenized using a teflon-glass homogenizer (15 strokes at 2000 rpm). Final concentration of 1 mg/ml was used for respiratory function assay. The mitochondrial respiration protocol included the following steps: (a) pyruvate 5mM + malate 2mM used to determine complex I (CI)-linked LEAK respiration; (b) ADP 5mM added to determine oxidative phosphorylation-dependent respiration (OXPHOS state); (c) glutamate 10mM added as an additional substrate for CI; (d) succinate 10mM, a complex II (CII) substrate, then added to reconstitute convergent CI- and CII-linked respiration; (e) rotenone (a C1 inhibitor) at 0.5μM end concentration added to measure CII-dependent respiration; and (f) antimycin A (a complex III inhibitor) at 2.5μM end concentration added to determine residual oxygen consumption, i.e. non-mitochondrial-dependent respiration. To determine complex IV-dependent respiration, N,N,N’,N’-tetramethyl-p-phenylenediamine dihydrochloride 0.5 mM and ascorbate 2 mM were added; the chemical background oxygen consumption was assessed after inhibiting CIV by using 100 mM azide. RCR was determined for each sample by dividing the OXPHOS-dependent respiration by LEAK respiration [from the above descriptions: (b)/(a)].

### Ischemic stroke model

ET-1 (a vasoconstrictive peptide) injection was used to induce transient middle cerebral artery occlusion in rats, which were anesthetized with 5% isoflurane in oxygen. At the start of surgery animals received SC tramadol (10mg/kg). Rats were placed in a stereotaxic apparatus (Stoelting Europe, Dublin, Ireland). Anesthesia was maintained with 2% isoflurane using a face mask. A midline incision was made in the skin above bregma, the skull cleaned from surrounding tissues, and a small hole drilled in the skull using a micromotor high-speed drill (Stoelting Europe). The stereotaxic coordinates for ET-1 microinjection (240pmol/rat or artificial cerebrospinal fluid for sham group) were 0.2mm anterior, 5.1mm lateral (to the left), and 8.2mm ventral relative to bregma, which is adjacent to the mid cerebral artery in the piriform cortex. 3.2μl of ET-1 (240pM) was injected at 1μl/min using a 10μl Hamilton syringe with 26s-gauge needle. Prior to slow withdrawal, the needle was left in the injection site for 3.5 minute to prevent back flow of ET1 or saline. The hole left after microinjection was left open, skin was closed using 4-0 silk threads (Sofsilk, Covidien). The sham-operated animals were subjected to the same surgical procedure except artificial cerebrospinal fluid was used for microinjection.

### Bioanalytical method for pharmacokinetic measurement of SBT-272

Blood was collected into K-EDTA tubes after cardiac puncture following euthanasia. Blood was spun using standard methods to collect plasma, which was stored at –80°C for subsequent analysis. After blood collection, animals were perfused transcardially with cold PBS solution for 5–8 minutes to remove all blood. For bioanalytical analysis, the rat hemi-brain and the whole mouse brains were harvested, collected in microtubes, and frozen in liquid nitrogen. These brains were homogenized using bead rupture (Omni Bead Ruptor 24), 96μl of homogenate transferred to low-binding microtubes and kept at –80°C until analysis. Drug concentration in mouse and rat plasma and brain homogenates was measured by liquid chromatography tandem mass spectrometry.

### Mouse 5-day chronic pharmacokinetic studies

Mice (*n*=4 per group) were dosed SC for 5 consecutive days with 1, 5, and 7.5mg/kg/day SBT-272 as pH-adjusted saline (pH~6.8) at dose volume of 10ml/kg. Eight hours after the final dose, animals were euthanized with regulated flow of CO_2_, blood was collected via cardiac puncture, and brains were collected after transcardial perfusion for 5 minutes with cold PBS. Drug concentrations were measured. Drug concentration in the blood 8 hours after dosing was below the lower limit of quantification. The ng/g value of drug in the brain is converted to nM assuming 1g of mouse brain was dissolved in 1ml of artificial cerebrospinal fluid.

### Motor cortex cultures

P3 motor cortices were isolated from WT-UeGFP and prp-hTDP-43^A315T^-UeGFP mice, dissected, dissociated, and cultured on glass coverslips (4×10^4^ cells per 18-mm diameter coverslip) (Fisherbrand) coated with poly-L-lysine 10mg/ml (Sigma). Dissociated motor cortex was cultured in SFM [0.034 mg/l bovine serum albumin, 1mM L-glutamine, 25U/ml penicillin, 0.025mg/mL streptomycin, 35mM glucose, and 0.5% B27 in Neurobasal-A Medium (Life Technologies)] in a humidified tissue culture incubator in the presence of 5% CO_2_ at 37°C, as previously described (Ozdinler and Macklis, 2006). SBT-272 (10nM, 100nM and 1μM), edaravone (10nM, 100nM and 1μM), and AMX0035 (1mM sodium phenylbutyrate + 100μM taurursodiol) were added to SFM at the start of culture. Cultures were fixed after 3 days *in vitro* (DIV). CSMN retained their eGFP expression in culture and they were distinguished among other cortical cells and neurons.

### Immunocytochemistry

Antibodies used were: anti-GFP (1:1000, Invitrogen; or 1:1000, Abcam), anti-CHCHD3 (1:200, Proteintech), anti-FLAG clone M2 (1:500, Sigma F1804, St Louis, MO), anti-GFAP (1:1000; Invitrogen, Thermo Fisher Scientific), and anti-Iba1 (1:500; Wako). Appropriate secondary fluorescent antibodies (1:500, AlexaFluor-conjugated, Invitrogen) were added to the blocking solution at room temperature for 2 hours in the dark. Nuclei were counterstained with 4’,6-diamidino-2-phenylindole.

### Quantification of CSMN health

CSMN health was quantitatively assessed by measuring changes in axon length and arborization complexity of neurites with and without treatment. Images taken with a 20X objective on an epifluorescent microscope (Nikon) and CSMN processes were traced using the Simple Neurite Tracer plugin from Fiji (ImageJ, NIH). The longest neurite was selected to measure the axon length in μm. The aggregation of the neurite tracings centered at the soma generates a profile available for Sholl analysis to quantify the number of intersections at 5-μm radius intervals for each neuron.

### Assessment of mitochondrial dynamics

Images were taken with a 20X objective on an epifluorescent microscope (Nikon). Mitochondria are labeled by CHCHD3 immunocytochemistry and CSMN were detected with eGFP expression. CSMN with mitochondria present only in the soma, along the axon, and in neurites were counted. Percent distribution of CSMN with mitochondria present only in the soma, the axon, and neurites were determined both for untreated and SBT-272-treated cases (*n*=3 mice in 3 independent experiments).

### Correlative light electron microscopy

Mixed cortical cultures from prp-hTDP-43^A315T^ mice were plated on gridded glass coverslips on petri dishes (MetTek) and cultured for 3DIV either in SFM with or without 100nM SBT-272. Correlative light EM was performed as described previously (Arai and Waguri, 2019). After fixing with 2% paraformaldehyde (PFA) and 0.5% glutaraldehyde for 10 minutes at room temperature, eGFP-expressing CSMN were identified under epifluorescent microscopy and their location on the grid was marked prior to EM analysis. Cells were further fixed in 2% PFA and 2% glutaraldehyde for 10 minutes at room temperature and 50 minutes at 4°C. Cells were washed with 0.12 M phosphate buffer pH 7.4, and then treated with 1% osmium tetroxide and 1.5% potassium ferrocyanide (Sigma) in 0.12M phosphate buffer pH 7.4. Cells were dehydrated by an ascending series of alcohol (50, 70, 80, 90, and 100%) followed by treatment with epoxy resin for 24 hours. The grids were mounted on resin blocks and cured at 65°C for 3 days. The blocks were trimmed to contain CSMN before proceeding to ultra-thin sectioning. Resin blocks were sectioned on a Leica Ultracut UC6 ultramicrotome (Leica Inc., Nussloch, Germany). Sections (70nm) were collected on 200 mesh copper-palladium grids. Ultra-thin sections were counterstained on a drop of UranyLess solution (Electron Microscopy Sciences, Hatfield, PA) and 0.2% lead citrate. Grids were examined on FEI Tecnai Spirit G2 TEM (FEI Company, Hillsboro, OR) and digital images were captured on an FEI Eagle camera.

For EM analysis, CSMN from three independent experiments were used. The total number of mitochondria included in the analyses, number of mitochondrial clumps per CSMN, number of cristae per mitochondrion, and percentage of mitochondria with cristae were quantified. Statistical analysis was performed using one-way ANOVA with GraphPad Prism software.

### Mitochondria polarity assay

P3 motor cortices were isolated from prp-hTDP-43^A315T^-UeGFP mice, dissected, dissociated, and digested with 100 U/ml papain enzyme (Worthington) and 100U/ml DNase enzyme (NEB) at 37°C for 20 minutes. After digestion, tissue was gently triturated and a single cell suspension was prepared, as described (Gautam et al., 2022; Genc et al., 2022a). Cells were cultured on poly-L-lysine (10mg/mL, Sigma) coated 100-mm plastic dishes (Fisherbrand). One dish was treated with 100nM SBT-272 prepared in SFM and another with SFM only. Cells were allowed to grow at 37°C for 72 hours. Cells were treated with 50nM TMRE (Abcam) prepared in SFM for 20 minutes and incubated at 37°C. Cells were scrapped off the dish and resuspended in 1ml SFM containing 50nM TMRE. Thereafter, cells were subjected to flow cytometry analysis using FACSymphony A5 High-Parameter SORP Cell Analyzer (BD Bioscience). Intensity of red TMRE signal was measured in eGFP^+^ neurons and percentage of TMRE accumulation was calculated in eGFP^+^ neurons for each experiment.

### *In vivo* administration of SBT-272

WT-UeGFP and prp-hTDP-43^A315T^-UeGFP mice were randomly assigned to different treatment groups. Mice received 1 or 5mg/kg/day SBT-272 or 0.9% saline vehicle (negative control) daily by intraperitoneal injection starting at P60. At P120, mice were deeply anesthetized and perfused. The brain was dissected out followed by post-fixation in 4% paraformaldehyde overnight and sectioned at 50μm using a Leica vibratome (Leica VT1000S, Leica Inc., Germany). Serial floating sections (300μm apart) were processed for GFP, GFAP, and Iba1 immunohistochemistry, as described (Genc et al., 2021; Jara et al., 2019).

### Imaging and data collection for *in vivo* study

A Nikon AXR confocal microscope (Nikon Inc., Melville, NY) was used to acquire low- and high-magnification images. EM grids were examined on FEI Tecnai Spirit G2 TEM (FEI Company) and digital images were captured on a FEI Eagle camera. EM imaging was performed at the Center for Advanced Microscopy/Nikon Imaging Center (CAM), Northwestern University Feinberg School of Medicine, Chicago.

### Statistical analyses

All analyses were performed using GraphPad Prism software and performed in a blinded fashion whereby the individual performing the analyses was blind to the genotype and treatment groups. The D’Agostino and Pearson normality test was performed on all datasets. Statistical differences between two groups were determined using either a parametric (Student’s *t*-test) or a non-parametric test (Mann-Whitney test), when appropriate. For comparison of two groups, an unpaired Student *t*-test with Welch’s correction was used. Statistical differences between more than two groups were determined by one-way ANOVA followed by the Tukey’s *post-hoc* multiple-comparison test. Šídák’s multiple comparisons test was used for comparison of Sholl data between two samples. Statistically significant differences were taken at *P*<0.05. Any other statistical test is as reported in the figures.

## RESULTS

### Identification and characterization of SBT-272

SBT-272 is a novel, first-in-class, brain-penetrant small molecule (Fig. 1, Supplementary table S1), which was discovered following the pharmacological success of elamipretide, a peptide that binds to cardiolipin and stabilizes the integrity of the IMM under mitochondrial stress (Allen et al., 2020; Mitchell et al., 2020; Szeto, 2014; Szeto, 2018; Szeto and Birk, 2014; Szeto and Liu, 2018). The activity of SBT-272 on mitigating mitochondrial dysfunction was assayed using well established *in vitro* and *in vivo* assays. The ability of SBT-272 to prevent the decline of mitochondrial respiratory function impacted by anoxia and reoxygenation was first determined *in vitro* in permeabilized rat cardiac fibers (Fig. 1A). Baseline respiration and overproduction of the reactive oxygen species H_2_O_2_ was measured with vehicle or 100nM SBT-272 treatment. Anoxia-reoxygenation had a statistically significant effect on reducing respiration and increasing H2O2 production compared to baseline in vehicle and SBT-272 treated wells (*P*<0.05). SBT-272 significantly reduced H_2_O_2_ production compared to the vehicle treatment (all values are mean ± SEM and expressed as a factor of each baseline (pre-anoxia) value, Vehicle: 1.6±0.1 and SBT-272: 1.3±0.1 for H_2_O_2_ production, *P*=0.023; Vehicle: 1.9±0.1 and SBT-272: 1.6±0.1 H_2_O_2_/O_2_ ratio, *P*=0.043). The effect of SBT-272 in preventing ischemia-reperfusion injury to mitochondria *in vivo* was measured in rats after an acute renal ischemia-reperfusion (Fig. 1B). SBT-272 significantly reduced the production of blood urea nitrogen (BUN) and creatinine, two established renal injury markers measured in the plasma (all values are mean ± SEM, BUN levels from vehicle, pre-ischemia: 11.4±0.5 and post-ischemia: 50.8±8.1mg/dL; SBT-272 at 4mg/kg, pre-ischemia: 11.6±0.9 and post-ischemia: 19.2±1.4mg/dL; *P*=0.002 for both compared to vehicle and pre-ischemia levels; creatinine levels: vehicle, pre-ischemia: 0.19±0.01 and post-ischemia: 1.5±0.4mg/dL; SBT-272 (4mg/kg), pre-ischemia: 0.2±0.0 and postischemia: 0.4±0.1mg/dL; *P*=0.017 & 0.008 compared to vehicle and pre-ischemia levels, respectively). SBT-272 exhibited optimized absorption, distribution, metabolism, and excretion (ADME) properties, such as high plasma half-life and biodistribution across the blood-brain-barrier, as evaluated in rats after a single dose of 5mg/kg delivered subcutaneously (SC) (Fig. 1C, Supplementary table S1).

**Fig. 1.**
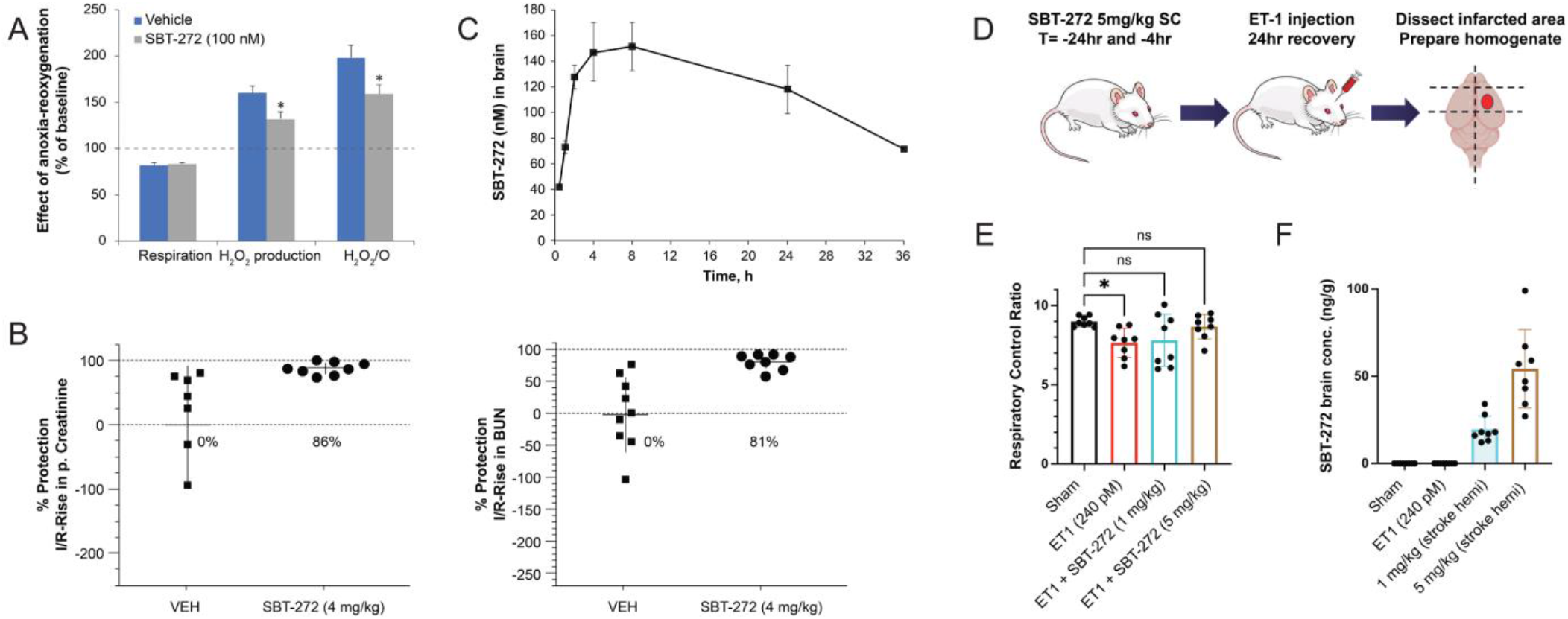
Pharmacodynamic effects of SBT-272 in rodent studies. (A) SBT-272 reduced mitochondrial H_2_O_2_ emission in rat cardiac fibers exposed to anoxia-reoxygenation stress (\**P*<0.05). (B) Plasma creatinine and blood urea nitrogen (BUN) levels were significantly decreased by SBT-272 4mg/kg SC *in vivo* in a rat after an acute kidney ischemia-reperfusion. (C) Rat brain SBT-272 concentrations up to 36 hours after a single dose of SBT-272 5mg/kg SC. (D) SBT-272 (5mg/kg SC, 4 and 24 hours before injury) prevented the loss of mitochondrial respiratory control ratio (RCR) in rat brain following cerebral ischemia-reperfusion injury induced via stereotactic delivery of 240pM ET-1 (artificial cerebrospinal fluid in sham) in the piriform region. Dotted lines indicate left and right subsection of hemibrain used for RCR and PK analysis. (E) Individual Respiratory Control Ratios (RCR) per animal (*n*=8 per group) from the stroke-induced subsection of the right hemisphere. (F) SBT-272 level in the brain homogenates from the same hemisphere after ET-1 injection. ET1 = endothelin-1, vasoconstrictive peptide (\**P*<0.05, Kruskal-Wallis followed by Dunn’s test).

Past biophysical and electron microscopy (EM) studies have demonstrated that elamipretide binds to mitochondrial cardiolipin and this can restore mitochondrial cristae structural defects that occur in various peripheral mitochondrial injury models (Allen et al., 2020). SBT-272 also partitions in the brain mitochondrial fraction (145.3nM in mitochondrial fraction compared to 66.5nM in brain homogenate) measured at 24 hours after administration of 5mg/kg SBT-272 SC treatment in rats (Allen et al., 2020). To assess its pharmacodynamic effect on mitochondrial function in the brain, SBT-272 was tested in a rat model of acute ischemia. The mitochondrial effects of SBT-272 were determined using the respiratory control ratio (RCR), the ratio of coupled respiration (supporting oxidative phosphorylation) to respiratory rate in the absence of ADP (‘leak’ respiration). The high mitochondrial RCR in sham-treated animals (9.0±0.1) confirmed that the mitochondria were healthy, as the majority of respiration accounted for replenishing cellular ATP levels (Fig. 1E). Consistent with mitochondrial injury, RCR declined significantly in this rat stroke model compared to controls (Fig. 1E). SBT-272 pretreatment 4 and 24 hours before endothelin toxin-1 (ET-1)-induced stroke injury protected against the stroke-induced reduction in mitochondrial RCR (Fig. 1E). The brain concentration 24 hours after final treatment was assessed in the same hemibrain used for RCR measurements of individual animals and was found to be dose proportional. The favorable pharmacology and *in vivo* biological activity in rodent brain motivated the exploration of SBT-272 especially within the context of TDP-43 pathology in ALS, as mitochondrial dysfunction emerges as one of the main and common underlying causes and pathologies of motor neuron vulnerability (Filosto et al., 2011; Gao et al., 2019; Gautam et al., 2019a; Gautam et al., 2019b; Hervias et al., 2006).

### SBT-272 improves mitochondrial integrity of CSMN diseased with TDP-43 pathology

UCHL1-eGFP mice, a reporter line for UMNs was previously generated (Yasvoina et al., 2013), in which CSMN (corticospinal motor neurons, also known as UMNs in mice) express eGFP, allowing precise visualization and investigation of CSMN at a cellular level both *in vivo* and *in vitro*. Prp-hTDP-43^A315T^-UeGFP mice were generated by crossbreeding UCHL1-eGFP (Fig. 2A, B) and prp-hTDP-43^A315T^ mice (Gautam et al., 2019a; Gautam et al., 2019b; Jara et al., 2019; Wegorzewska et al., 2009), a well-characterized mouse model of ALS with TDP-43 pathology (Fig. 2A, C). In these mice, CSMN that are diseased due to TDP-43 pathology express eGFP *in vivo* (Fig. 2D–G) and *in vitro* (Fig. 2H, I). TDP-43 pathology is present in diseased CSMN both *in vivo* (Fig. 2E,G) and even at 3 days *in vitro* (3DIV; Fig. 2I). Therefore, this model has unique advantages and strengths to visualize CSMN among many other cortical neurons and cells with cellular precision, and to investigate their cellular responses to compound/drug treatment.

**Fig. 2.**
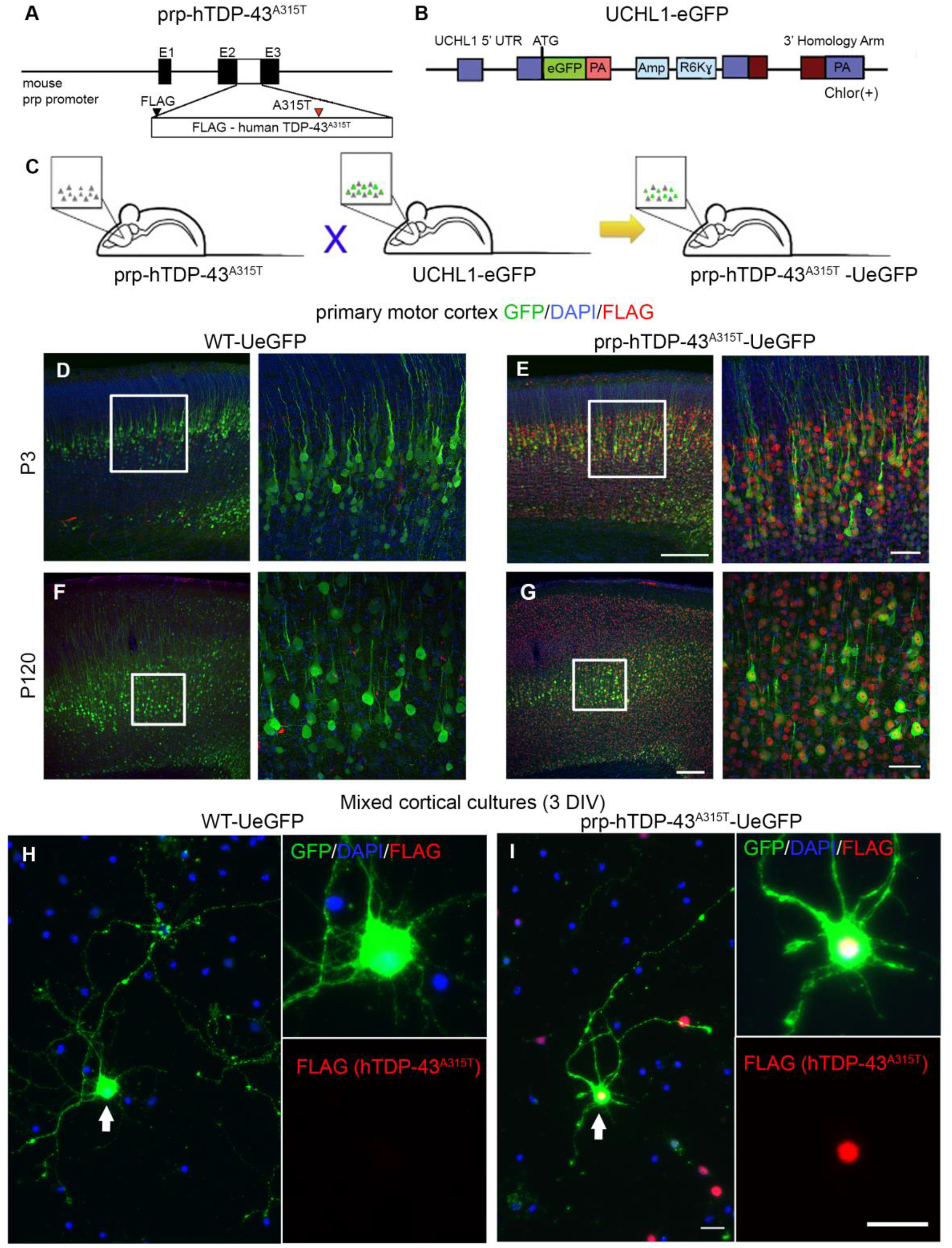
prp-hTDP-43^A315T^-UeGFP mice allow visualization and cellular assessment of CSMN with TDP-43 pathology both *in vivo* and *in vitro*. (A) Prp-hTDP-43^A315T^ mice express human TDP-43 protein with the A315T point mutation and an N-terminal FLAG tag. (B) Genetic construct of UCHL1-eGFP mice (C) Prp-hTDP-43^A315T^ are crossed with the UCHL1-eGFP to generate prp-hTDP-43^A315T^-UeGFP mice, CSMN reporter lines with TDP-43 pathology. (D, E) P3 motor cortex image of WT-UeGFP (D) and prp-hTDP-43^A315T^-UeGFP (E) mice. (F,G) P120 motor cortex image of WT-UeGFP (F) and prp-hTDP-43^A315T^-UeGFP (G) mice. CSMN are in layer 5 of the motor cortex. FLAG staining (red dots) help visualize the N-terminus FLAG tag attached to the exogenous human TDP-43^A315T^ protein in the nuclei of neurons, including CSMN of prp-hTDP-43^A315T^ mice. Scale bars: 200μm in low magnification (left) and 50μm in high magnification (right) panels. (H, I) Representative images of mixed cortical cultures isolated from the motor cortex of WT-UeGFP (H) and prp-hTDP-43^A315T^-UeGFP (I) mice. CSMN retain their eGFP expression *in vitro* (arrow). Diseased CSMN have FLAG tag of hTDP-43^A315T^. (blue=DAPI stain; red=FLAG tag). Scale bars: 20μm.

Since EM helped visualize and reveal intracellular defects that occur in CSMN with high-level precision, we performed EM to visualize potential improvements that are mediated by 100 nM SBT-272 treatment [CSMN (*n*=3) from independent experiments (*n*=3) were used (Fig. 3)]. The choice of 100nM is justified by observed potency reported in the rat cardiac cells. eGFP-expressing CSMN were identified under an epifluorescent microscope, marked and processed for EM analysis (Fig. 3A,A’), which revealed the integrity of IMM in CSMN of WT-UeGFP (Fig. 3B, B’,B’’), of untreated prp-hTDP-43^A315T^-UeGFP mice (Fig. 3C,C’,C’’) and SBT-272 treated prp-hTDP-43^A315T^-UeGFP (Fig. 3D,D’,D’’) mice. The total number of mitochondria, number of mitochondrial clumps, number of cristae per mitochondrion, and the percentage of mitochondria with cristae were quantified and statistical analyses were performed using unpaired Student *t*-tests with GraphPad Prism software (Supplementary table S2).

**Fig. 3.**
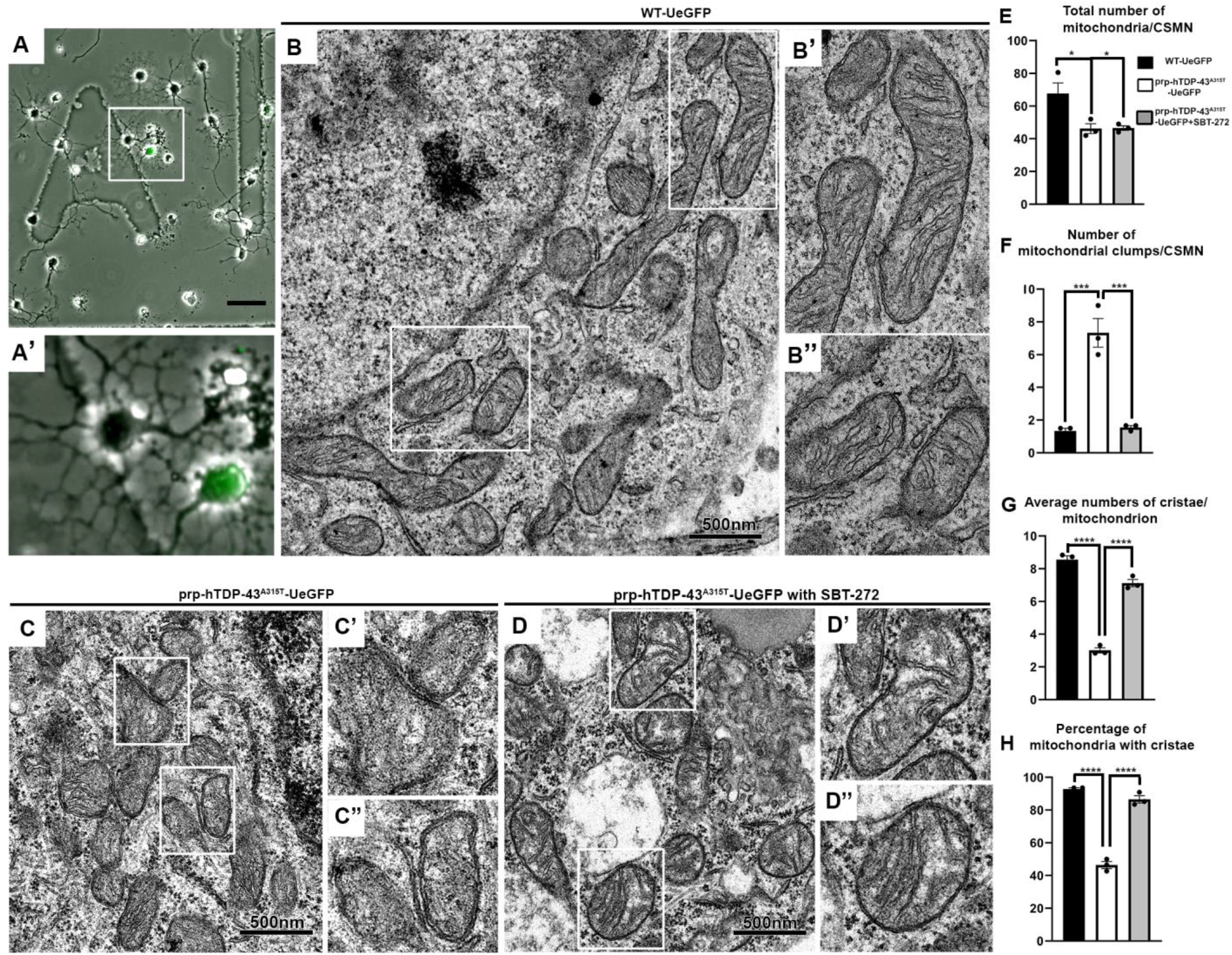
SBT-272 treatment improves ultrastructural integrity of mitochondria in diseased CSMN. (A, A’) CSMN of prp-hTDP-43^A315T^-UeGFP mice express eGFP. Their location on the grid is marked prior to EM analysis. (B–B”) Representative EM images of mitochondria inside CSMN of WT-UeGFP mice. (C–D”) Representative EM images of mitochondria inside CSMN of prp-hTDP-43^A315T^-UeGFP mice untreated (C–C”) or treated with 100nM SBT-272 (D–D”). Scale bars: 50μm in (A) and 500nm in (B-D). (E–H) Quantitative assessment of mitochondrial health upon SBT-272 treatment. Quantification of the total number of mitochondria/CSMN (E), number of mitochondrial clumps per CSMN (F), average numbers of cristae per mitochondrion (G), and percentage of mitochondria with cristae in prp-hTDP-43^A315T^-UeGFP mice untreated or treated with 100nM SBT-272 (H). Mean±SEM numbers are shown for each group. \**P*<0.05, \*\**P*<0.01, \*\*\**P*<0.001, one-way ANOVA with Tukey’s multiple comparisons test.

The average number of mitochondria per CSMN did not change after SBT-272 treatment (prp-hTDP-43^A315T^-UeGFP:46.2±2.9; prp-hTDP-43^A315T^-UeGFP+100nM SBT-27: 46.5±1.4; adjusted *P*=0.998, Fig. 3E), suggesting that SBT-272 treatment did not promote mitochondrial proliferation. However, the integrity and stability of mitochondria improved upon SBT-272 treatment. CSMN in wild type (WT) healthy control mice contained a significantly higher percentage of healthy and intact mitochondria compared to CSMN that are diseased with TDP-43 pathology (WT-UeGFP: 67.7±6.4%; prp-hTDP-43^A315T^-UeGFP: 46±2.9%, adjusted *P*=0.0259, Fig. 3E). In some cases, three or more mitochondria were found to aggregate. They were considered “clumps of mitochondria” and were quantified. Upon SBT-272 treatment, the number of mitochondrial clumps per CSMN significantly decreased in prp-hTDP-43^A315T^ mice (SFM: 7.3± 0.8; SBT-272: 1.5±0.1, adjusted *P*=0.0006), and were comparable to that of WT CSMN (WT-UeGFP: 1.3±0.2, adjusted *P*=0.954, Fig. 3F). Furthermore, the mean (±SEM) number of cristae per mitochondrion in CSMN of prp-hTDP-43^A315T^ mice significantly increased with SBT-272 treatment (SFM: 3.01±0.5; SBT-272: 7±0.2, adjusted *P*=0.0001, Fig. 3G) and were comparable to the number of cristae per mitochondrion in CSMN of healthy mice (WT-UeGFP: 8.6±0.2, adjusted *P*=0.005). The mean percentage of mitochondria with intact cristae were high in CSMN of WT neurons (WT-UeGFP: 92.8±0.7, adjusted *P*=0.110, Fig. 3H) and the percentage of mitochondria with intact cristae also significantly increased in the CSMN of prp-hTDP-43^A315T^ mice upon SBT-272 treatment (SFM: 46.4±2.1%; SBT-272: 86.5± 2.2%, adjusted *P*=0.0001).

### SBT-272 increases mitochondrial membrane polarization in diseased UMNs

We next investigated whether structural amendment of disrupted IMM leads to functional improvement. IMM integrity is crucial for maintaining membrane potential, which is required for normal mitochondria function (Duchen, 1999). Thus, assessment and quantification of mitochondrial membrane polarization had been a measure of mitochondrial function and health (Kuhlbrandt, 2015; Lin and Sheng, 2015; Miller and Sheetz, 2004; Paumard et al., 2002; Verburg and Hollenbeck, 2008). The fluorescent lipophilic cationic dye, tetramethylrhodamine ethyl ester (TMRE) (O’Reilly et al., 2003) rapidly accumulates in polarized mitochondria and depolarization leads to loss of TMRE fluorescence intensity, allowing for quantitation of membrane polarization and the extent of IMM integrity. Dissociated cell cultures were prepared from the motor cortex of prp-hTDP-43^A315T^-UeGFP mice at postnatal day 3 (P3) and treated with SBT-272 (100nM) for 72 hours prior to flow cytometry analysis. Untreated mice were used as negative control. Forward-scatter and side-scatter plots were generated, and gates were determined first to exclude clumps and debris, then to select only live cells and, finally, to select and investigate CSMN (i.e GFP^+^ neurons among live cells). The TMRE levels (extent of red fluorescence) were measured in CSMN of prp-hTDP-43^A315T^-UeGFP mice that were untreated (Fig. 4A) and treated with SBT-272 (Fig. 4B). The TMRE levels of untreated CSMN were taken as baseline (i.e. 100%) and relative percent change of TMRE levels in CSMN of prp-hTDP-43^A315T^-UeGFP mice treated with SBT-272 were determined for all three independent experiments (marked by “*”, “ •” and “∆”; Fig. 4C). TMRE levels were significantly higher upon SBT-272 treatment in all three independent experiments (prp-hTDP-43^A315T^: 100±0%; prp-hTDP-43^A315T^+SBT-272: 142±15%, *n*=3, *P*=0.04, Fig. 4C), suggesting that SBT-272 treatment also improved mitochondrial membrane potential and function.

**Fig. 4.**
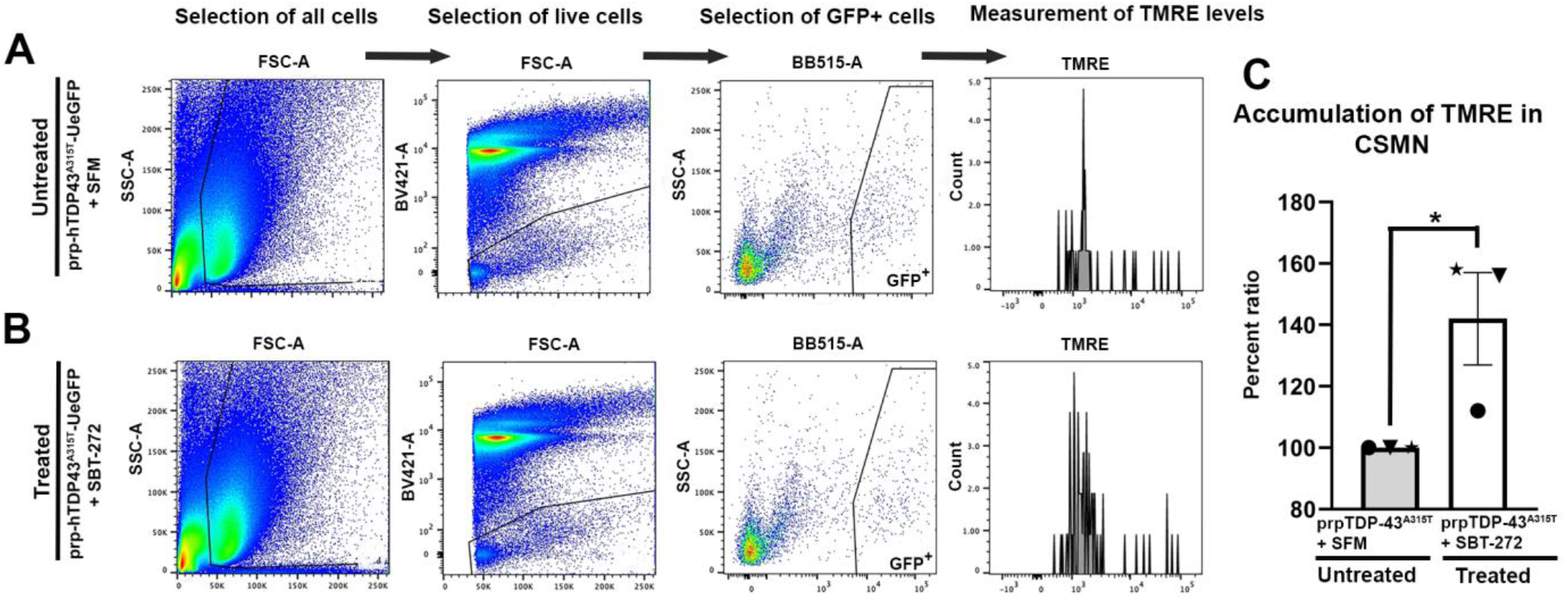
SBT-272 improves mitochondria membrane polarization. (A,B) Flow-sorting and gating strategy for selecting all cells (first column), live cells (second column), CSMN (eGFP^+^ cells; third column), and the extent of TMRE accumulation in CSMN (fourth column) using dissociated cortical cultures isolated from prp-hTDP-43^A315T^-UeGFP mice and that are treated with SFM (A) and 100nM SBT-272 (B). (C) Bar graph representation of TMRE levels in treated and untreated cases in three independent experiments (“*”, “ •” and “∆”). The untreated cases are marked as 100%, and the relative percent TMRE accumulation are determined in treated cases, for each experiment. Mean ± SEM is shown for each treatment group. \**P*<0.05, Student’s *t*-test.

### SBT-272 treatment improves mitochondrial motility in diseased CSMNs

Because improved IMM integrity leads to improved axonal transport of mitochondria (Lin and Sheng, 2015; Miller and Sheetz, 2004; Verburg and Hollenbeck, 2008), and since their detection along the axon of CSMN is a clear indication of the neuron’s ability to move them along the newly formed axon, we next investigated whether SBT-272 treatment also enhances mitochondrial transport along the axon. In diseased CSMN, mitochondria were mainly restricted to the soma (Fig. 5 A), and only 18.8±4% of CSMN had mitochondria present along the axon (Fig. 5C). In contrast, 75±5% of healthy CSMN had mitochondria present both in the soma and in the axon. SBT-272 treatment dose-dependently increased the percentage of diseased CSMN that harbor mitochondria along their axon and neurites, compared to diseased CSMN cultured in SFM (10nM SBT-272: 36.7±4% of CSMN, adjusted *P*=0.0912; 100nM SBT-272: 53.2±5% CSMN, adjusted *P*=0.0017; 1μM of SBT-272: 63.3±4%, adjusted *P*=0.0002).

**Fig. 5.**
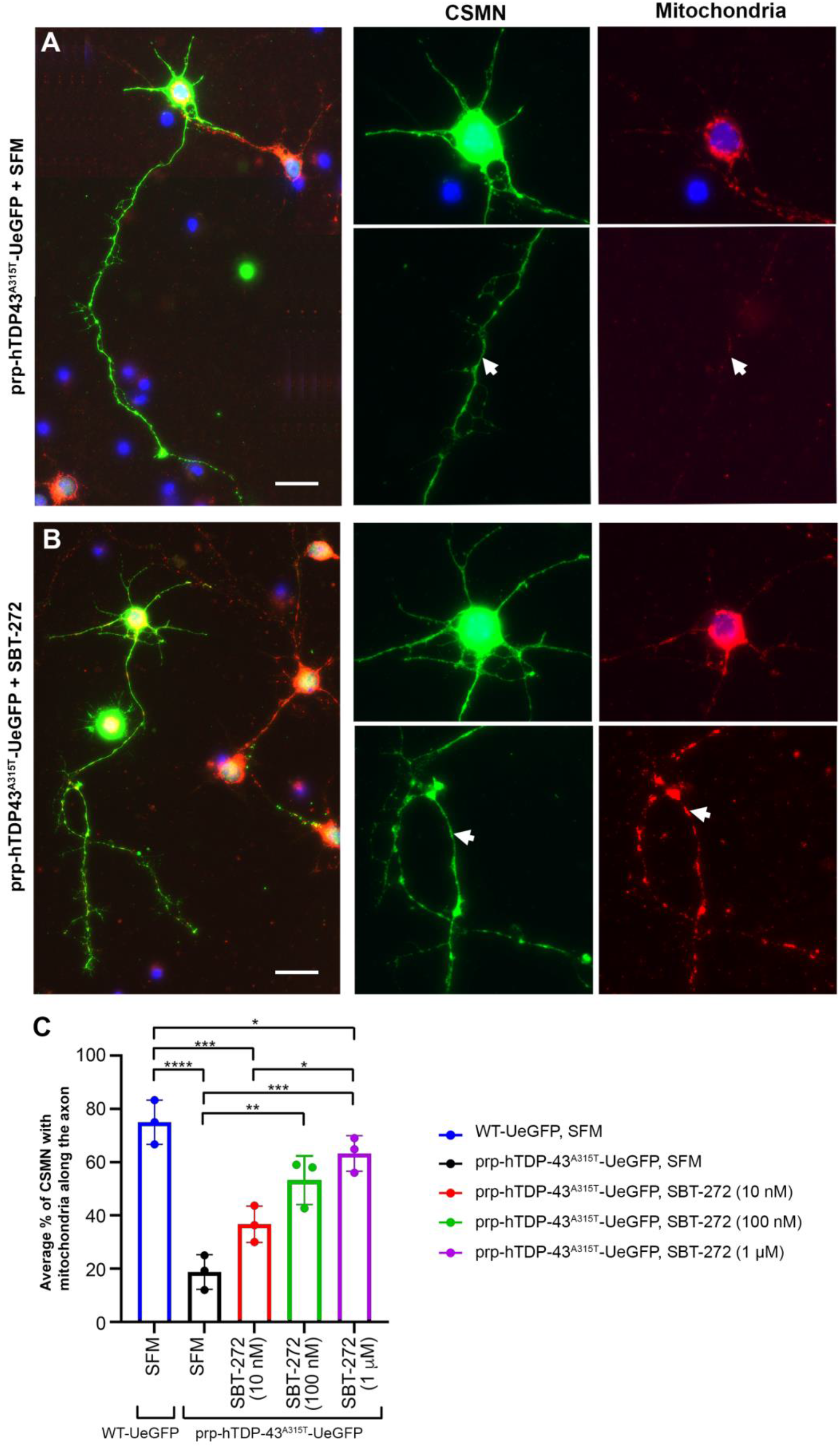
SBT-272 treatment improves mitochondrial motility of diseased CSMN. (A) Mitochondria are restricted to the soma in diseased CSMN that are cultured in SFM, and are not detected along the axon (white arrow). (B) Upon SBT-272 treatment, mitochondria become detected in the axon and in neurites of diseased CSMN (white arrow). (C) Bar graph representation of percent CSMN with mitochondria detected in axon after different doses of SBT-272 treatment (10nM, 100nM, 1μM) and with SFM treatment. CHCHD3 immunocytochemistry is used to visualize mitochondria. Each dot represents the mean value per mouse. Three independent mice were used for each treatment and genotype. Mean±SEM is shown for each treatment group. \**P*<0.05, \*\**P*<0.01, \*\*\**P*<0.001, \*\*\*\**P*<0.0001, one-way ANOVA with Tukey’s multiple comparisons test. Scale bars: 50μm.

### SBT-272 treatment improves CSMN health *in vitro*

We next investigated whether improvement of IMM structure, mitochondrial membrane function, and mitochondrial motility lead to improved UMN health at a cellular level. When dissociated cortical cultures are established, CSMN are distinguished among many different cortical cells and neurons due to their eGFP expression, UMN morphology with prominent apical dendrite, large pyramidal body, and a long axon (Dervishi and Ozdinler, 2018). In addition, they express CTIP2, a molecular marker specific to CSMN (Dervishi and Ozdinler, 2018). We previously established CSMN cultures using motor cortex isolated from UCHL1-eGFP and prp-hTDP-43^A315T^-UeGFP mice at P3 (Ozdinler and Macklis, 2006). In these cultures, CSMN of UCHL1-eGFP mice appeared healthy with a large pyramidal cell body, an apical dendrite, and a well-defined axon. In contrast, CSMN of prp-hTDP-43^A315T^-UeGFP mice displayed the morphology of a diseased neuron (Ozdinler and Macklis, 2006). For example, average axon length was significantly shorter (WT: 371.7±10.9μm, *n*=3; prp-hTDP-43^A315T^: 154.7±5.9μm, *n*=3, *P*<0.0001) and the majority of diseased CSMN that had shorter axons (Gautam et al., 2022). They also displayed significantly less branching and arborization. We, therefore, utilized average axon length and the extent of branching and arborization as two quantitative outcome measure to assess the cellular responses of diseased CSMN to compound treatment (Al-Ali et al., 2016; Gautam et al., 2022; Genc et al., 2022a; Nguyen et al., 2012; Ozdinler and Macklis, 2006; Ristanovic et al., 2006; Sholl, 1953; Whitlon, 2017).

While we investigated the activity of SBT-272 to promote and enhance axonal outgrowth of diseased CSMN, we chose to evaluate two additional compounds known to act on cellular stress and have been investigated in preclinical models of ALS and in patients. These compounds are edaravone, FDA approved for ALS (Brooks et al., 2022; Genge et al., 2022; Ito et al., 2008; Soejima-Kusunoki et al., 2022; Writing and Edaravone, 2017; Yoshino and Kimura, 2006), and AMX0035, an investigational compound approved for the treatment of ALS in Canada (Heo, 2022). Edaravone is believed to be a scavenger of free radical such as 3-nitrotyrosine that cause cellular stress (Ikeda and Iwasaki, 2015; Yoshida et al., 2008; Yoshino and Kimura, 2006) and AMX0035 is reported to be effective in improving the health and stability of mitochondria and endoplasmic reticulum (Paganoni and Cudkowicz, 2020; Paganoni et al., 2022a; Paganoni et al., 2020; Paganoni et al., 2022b).

A dose-response curve helped determine the optimal concentration of SBT-272 necessary to improve CSMN axon length (Fig. 6, Supplementary table S3). Mixed motor cortex cultures were prepared from prp-hTDP-43^A315T^-UeGFP mice (Gautam et al., 2019a; Yasvoina et al., 2013). Approximately 40,000 cells were plated per well and were kept in culture for 3 days *in vitro* (3DIV) in the presence of either SFM (negative control) or with the addition of SBT-272 (10nM, 100nM, or 1μM). CSMN of prp-hTDP-43^A315T^-UeGFP mice had an average axon length of 154.7±5.9 μm when cultured in SFM (Fig. 6A, G). However, SBT-272 treatment significantly increased axon outgrowth (10nM SBT-272: 360.1±20.6μm, adjusted *P*<0.0001; 100nM SBT-272: 418.9±25.9μm, adjusted *P*<0.0001; 1μM SBT-272: 291.6±23.4μm, adjusted *P*=0.0029, Fig. 6B–D,G), suggesting that the optimal dose of SBT-272 treatment is at 100nM.

**Fig. 6.**
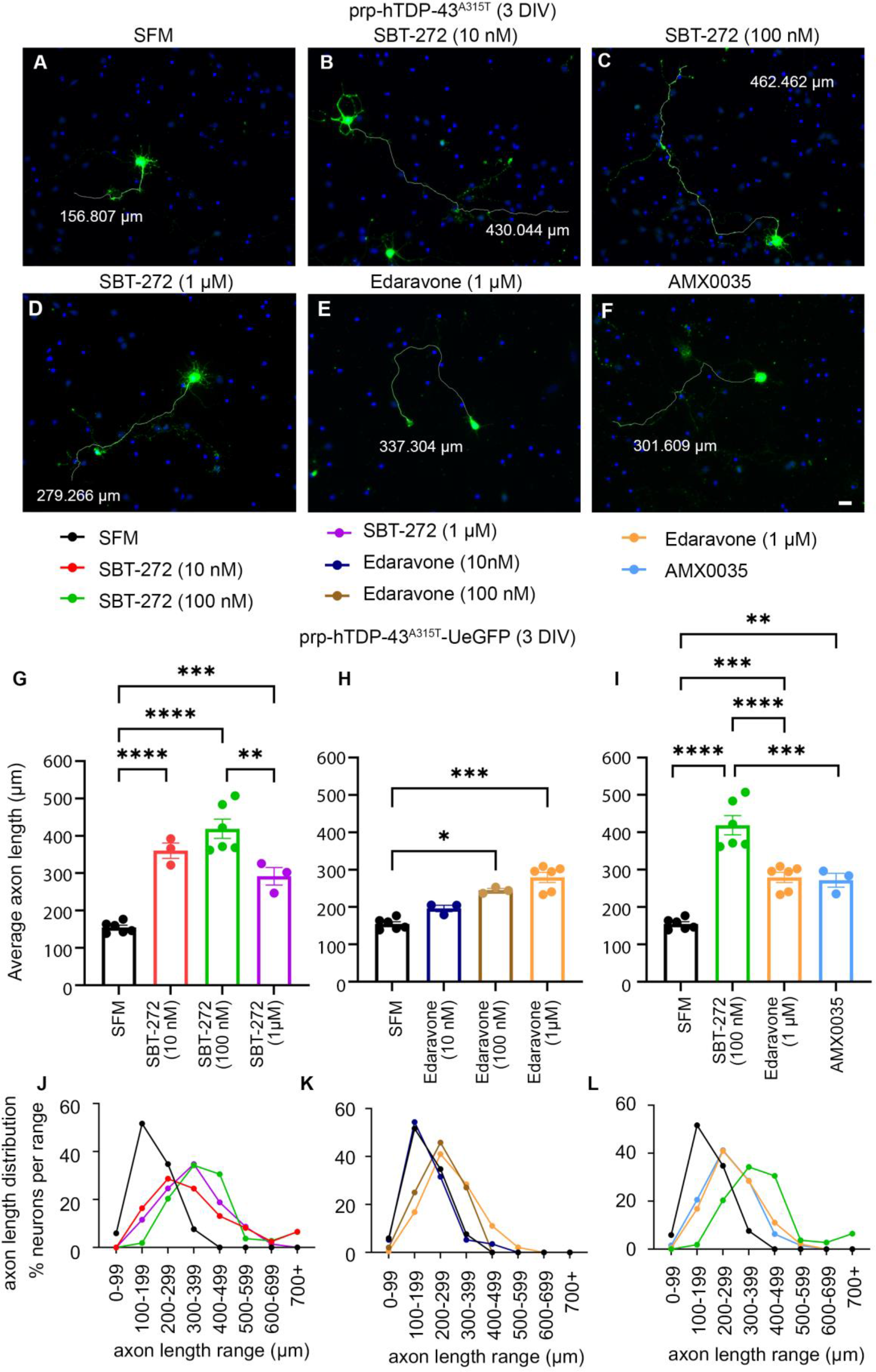
SBT-272 enhances axon outgrowth of diseased CSMN. (A–F) Representative images of CSMN from prp-hTDP-43^A315T^-UeGFP mice with various drug treatments after 3DIV. Blue dots (DAPI) represent other cells in culture, whereas CSMN are identified by their eGFP expression. (A) Untreated control CSMN grown in SFM. (B–D) CSMN of prp-hTDP-43^A315T^-UeGFP mice treated with 10 nM (B), 100nM (C), and 1μM (D) of SBT-272. (E–F) CSMN of prp-hTDP-43^A315T^-UeGFP mice treated with 1μM edaravone (E) and AMX0035 (F). Scale bar: 20μm. (G–I) Quantification of longest axon length of CSMN of prp-hTDP-43^A315T^-UeGFP mice with the aforementioned treatments. Each dot represents an independent mouse and each bar graph is color-coded. Mean±SEM length of the longest axon is shown for each treatment group. \**P*<0.05, \*\**P*<0.01, \*\*\**P*<0.001, \*\*\*\**P*<0.0001; one-way ANOVA with Tukey’s multiple comparisons test. (J–L) Percent distribution of axon length of diseased CSMN upon distinct compound treatment (color coded lines match colors of bar graph).

We next determined the dose-response curve for edaravone. At 10nM, average CSMN axon length was similar to the SFM-treated cases (196.3±8.6μm), whereas 100nM and 1μM concentrations displayed a significant dose-dependent increase (100nM edaravone: 244.8±4.7μm, adjusted *P*=0.0003; 1μM edaravone: 279.2±13.6 μm, adjusted *P*< 0.0001, Fig. 6E, H), suggesting that 1μM concentration would be most feasible to test *in vitro*.

As the optimal *in vitro* concentration for AMX0035 has been reported to be 1mM sodium phenylbutyrate+100μM tauroursodiol, we tested and compared all three compounds at their optimal concentrations for their ability to enhance axon outgrowth of diseased CSMN (Fig. 6F,I). Even though edaravone and AMX0035 significantly improved CSMN axon outgrowth (1μM edaravone: 279.2±13.6μm; AMX0035: 271.7±18.7μm), SBT-272 was more effective and was significantly better than both edaravone (SFM versus 100nM SBT-272, adjusted *P*<0.0001; SFM versus 1μM edaravone, adjusted *P*=0.0004; 100nM SBT-272 versus 1μM edaravone, adjusted *P*<0.0001) and AMX0035 (SFM versus AMX0035, adjusted *P*=0.0045; 100nM SBT-272 versus AMX0035, adjusted *P*=0.0005).

Percent axon length distribution analyses also revealed a robust enhancement of axon outgrowth upon SBT-272 treatment. SBT-272 (100nM) effectively shifted the percent distribution of axon length (Fig. 6J). Edaravone at 10nM was similar to SFM treatment, whereas 100nM and 1μM doses improved axon length distribution (Fig. 6K). When SBT-272, edaravone, and AMX0035 were compared, SBT-272’s ability to promote axon outgrowth became more evident. More than half of CSMN treated with 100nM SBT-272 had axons >300μm in length (Fig. 6L), which was not observed with edaravone or AMX0035 treatment.

The extent of branching and arborization that dissociated neurons manifest is related to axonal outgrowth but also a more robust measure of their health (Clark et al., 2016; Ristanovic et al., 2006). For example, CSMN of prp-hTDP-43^A315T^-UeGFP mice had significantly less branching and arborization. However, when diseased CSMN were treated with SBT-272 (10 and 100nM), their ability to branch and arborize was significantly improved (Fig. 7; all statistical analyses are reported in Supplementary table S4). SBT-272 (10 and 100nM) was most effective in increasing the number of intersections between the 50–200 μm range from the cell body and were indistinguishable in the >100 μm range. The number of intersections with the highest dose of SBT-272 (1μM) was significantly smaller. However, the optimal dose of 100nM, as previously determined by axon length (Fig. 6G), also displayed enhanced branching and arborization, especially in the range of 65–235μm from the cell body, further suggesting the optimal dose for SBT-272 is 100nM *in vitro* (Fig. 7A–D, G). When edaravone (Fig. 7E, H) and AMX0035 (Fig. 7F, I) were compared to SBT-272 for their ability to promote branching and arborization at their optimal concentrations, AMX0035 was more effective in enhancing branching and arborization of somatic neurites, whereas SBT-272 was able to promote much more effective branching and arborization, especially along the axon (Fig. 7I, Supplementary table S4).

**Fig. 7.**
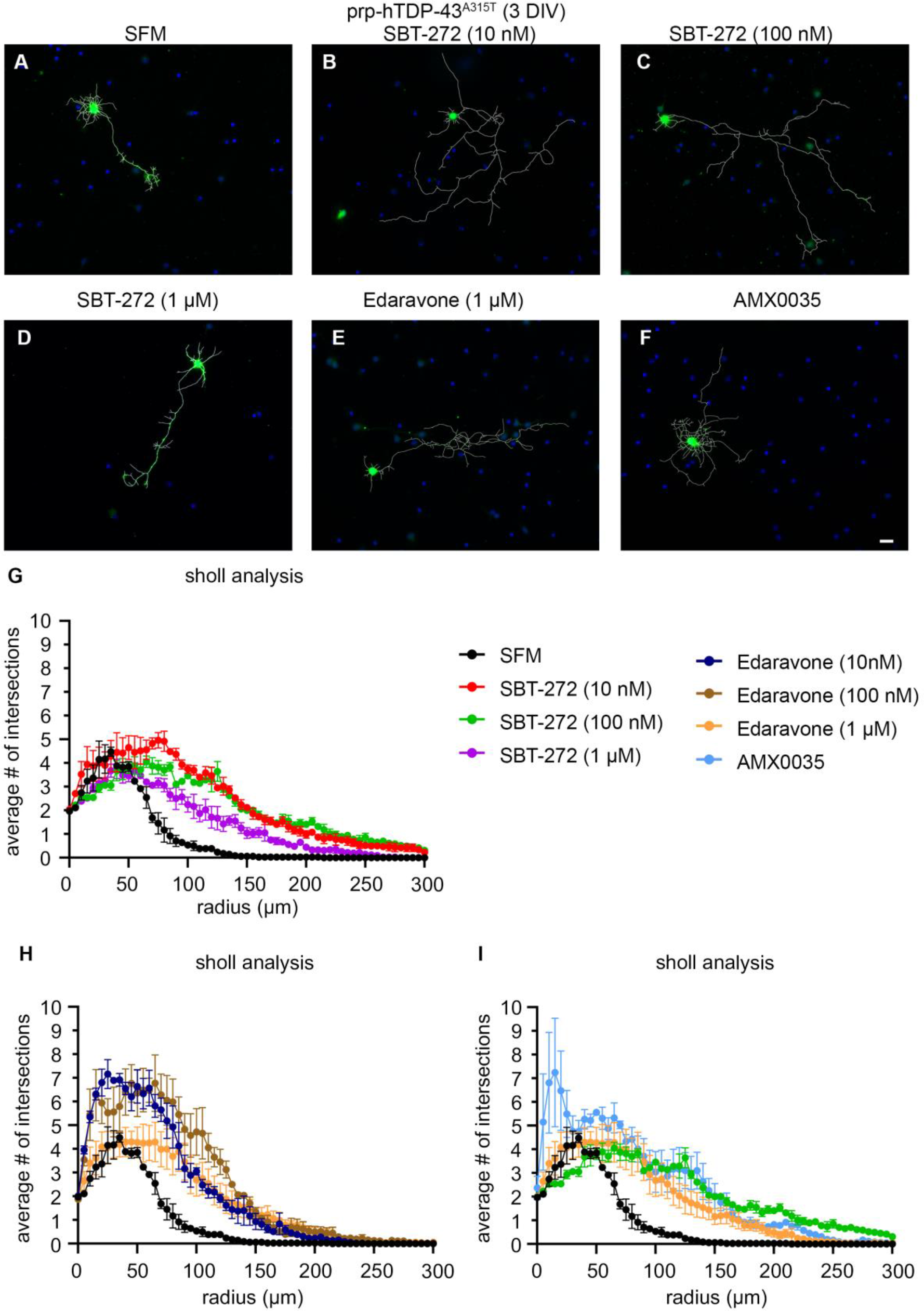
SBT-272 enhances branching and arborization of diseased CSMN. (A–F) Representative images of CSMN from prp-hTDP-43^A315T^-UeGFP mice with various drug treatments for 3DIV. Blue dots (DAPI stain) represent nuclei of all cells in culture, whereas CSMN are identified by eGFP expression. (A) Untreated control CSMN grown in SFM only. (B–D) CSMN of prp-hTDP-43^A315T^-UeGFP mice treated with 10nM (B), 100nM (C), and 1μM (D) of SBT-272. (E–F) CSMN of prp-hTDP-43^A315T^-UeGFP mice treated with 1μM edaravone (E) and AMX0035 (F). Scale bar: 20μm. (G–I) Sholl analysis of CSMN from prp-hTDP-43^A315T^-UeGFP mice with or without the aforementioned drug treatments. Each treatment is color coded.

### SBT-272 provides neuroprotection to CSMNs diseased with TDP-43 pathology

We next investigated whether SBT-272 treatment would also have cellular benefits for diseased UMNs *in vivo* (Fig. 8). To investigate potential changes in CSMN numbers and the extent of astrogliosis and microgliosis in the motor cortex, prp-hTDP-43^A315T^-UeGFP mice were treated with vehicle (negative control) or SBT-272 via intraperitoneal injection starting at P60 till P120 (Fig. 8A).

**Fig. 8.**
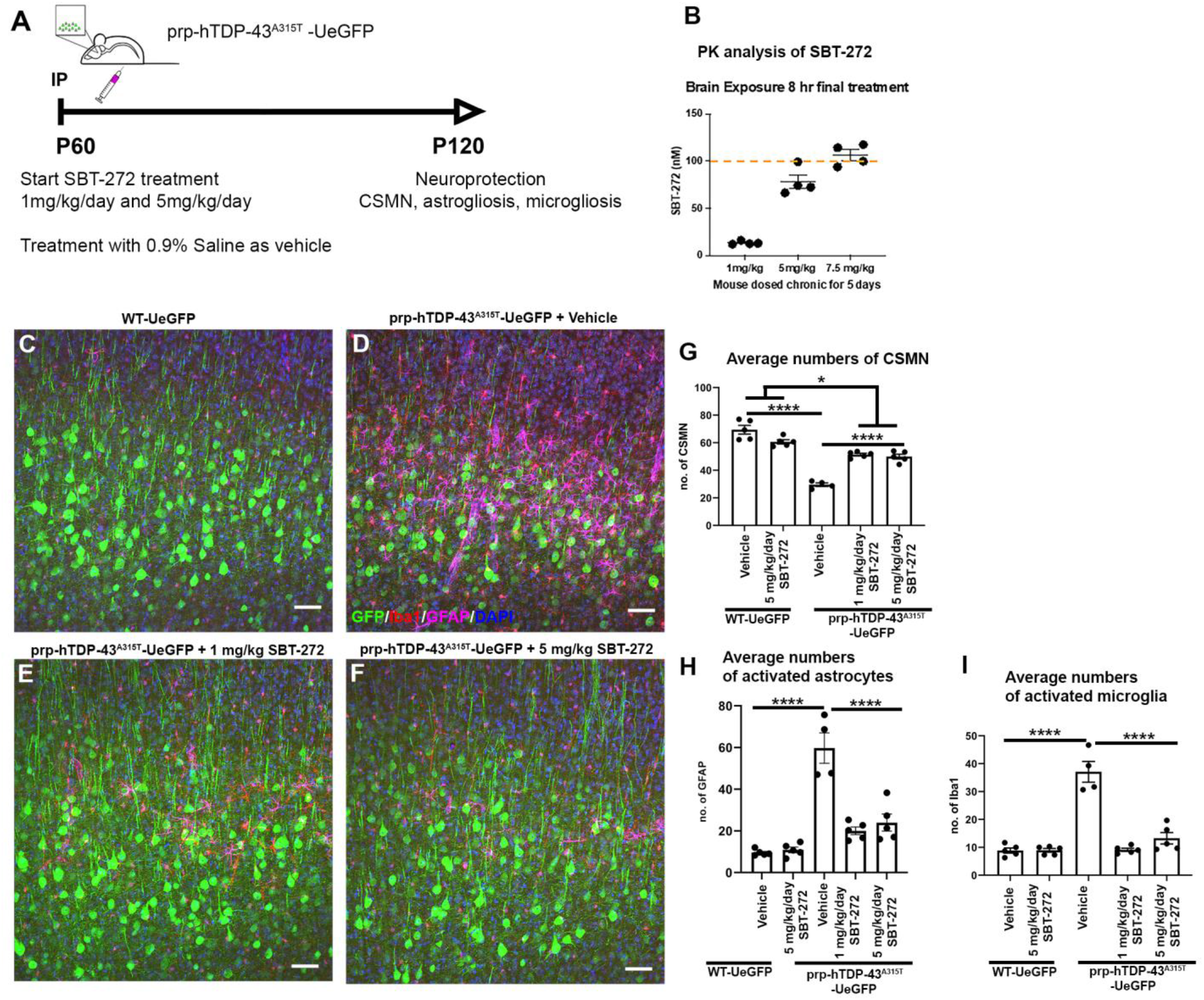
SBT-272 treatment is neuroprotective in the motor cortex of ALS with TDP-43 pathology. (A) Experimental design. (B) Pharmacokinetic analysis of SBT-272 brain exposure after 8 hours treatment. (C–F) Representative images of motor cortex of untreated WT-UeGFP mice (C), prp-hTDP-43^A315T^-UeGFP treated with vehicle (D), with 1mg/kg SBT-272 (E), and with 5mg/kg SBT-272 (F). Scale bars: 50μm. (G–I) Bar graph representation of CSMN numbers (G), average numbers of activated astrocytes (H), and activated microglia (I) in both WT-UeGFP and prp-hTDP-43^A315T^-UeGFP mice. Each dot represents the average value per mouse. Mean±SEM is shown for each treatment group. \**P*<0.05, \*\*\*\**P*<0.0001, one-way ANOVA with Tukey’s multiple comparisons test.

In lieu of targeting appropriate brain concentration during chronic dosing in mice to meet the *in vitro* potency range of 10–100nM, a pharmacology study was conducted in WT mice. Animals were dosed with SBT-272 1, 5, and 7.5mg/kg/day SC for 5 consecutive days to achieve steady-state brain concentrations. Blood and brains were harvested after 8 hours, i.e., the earliest time when the drug level in mouse brain is close to sustained trough levels as from a single dose pharmacokinetic study. All doses were well tolerated by the WT mice. The plasma concentration at 8 hours was below the limits of detection, as expected from the single dose pharmacokinetic study. Brain SBT-272 concentration was dose proportional with both 1 and 5mg/kg achieving mean trough concentrations of 10–75nM (*n*=4 mice/group) (Fig. 8B). Based on these pharmacokinetic calculations and assessments, prp-hTDP-43^A315T^-UeGFP mice were treated with two different doses of SBT-272 (1 and 5mg/kg/day).

The CSMN of prp-hTDP-43^A315T^-UeGFP mice were labeled with stable eGFP expression (Fig. 2). Since we recently found that the numbers of activated astrocytes and microglia are increased in the motor cortex of in prp-hTDP-43^A315T^ mice (Jara et al., 2019) and there is astrogliosis and microgliosis in the motor cortex of ALS patients with TDP-43 pathology (Jara et al., 2019), we quantified the extent of astrogliosis and microgliosis based on their cellular morphology and the markers they express in this study (Fig. 8C–F, Supplementary fig. S1).

CSMN numbers in WT mice did not change with or without SBT-272 treatment (WT-UeGFP+vehicle: 69.5±3.1, *n*=5; WT-UeGFP + 5mg/kg/day SBT-272: 60.7±1.5, *n*=5, Fig. 8G), suggesting good tolerability and no non-specific activity of chronic treatment with SBT-272. Since CSMN undergo progressive degeneration in prp-hTDP-43^A315T^-UeGFP mice, their numbers were reduced compared to WT-UeGFP mice (prp-hTDP-43^A315T^-UeGFP +vehicle: 29.6±1.3, *n*=4, versus WT-UeGFP+vehicle: 69.5±3.1, adjusted *P*<0.0001, Fig. 8G). These findings agree with the timing and the extent of CSMN loss in prp-hTDP-43^A315T^ mice, as previously reported (Gautam et al., 2019a). Upon SBT-272 treatment, however, there were significantly more CSMN retained in the motor cortex of prp-hTDP-43^A315T^-UeGFP mice (prp-hTDP-43^A315T^-UeGFP+1mg/kg/day SBT-272: 51.4±0.9, *n*=5; prp-hTDP-43^A315T^-UeGFP+5 mg/kg/day SBT-272: 49.9±1.8, *n*=5, adjusted *P*<0.0001, Fig. 8G).

### SBT-272 treatment reduces astrogliosis and microgliosis in the motor cortex *in vivo*

There were significantly higher numbers of activated astrocytes in the motor cortex of prp-hTDP-43^A315T^-UeGFP mice compared to that of WT-UeGFP mice (prp-hTDP-43^A315T^-UeGFP+vehicle: 59.8 ± 7.3, *n*=4, versus WT-UeGFP+vehicle: 9.5±0.7, *n*=5, adjusted *P*< 0.0001, Fig. 8H). The number of activated astrocytes was reduced upon SBT-272 treatment at both doses (prp-hTDP-43^A315T^-UeGFP+5 mg/kg/day SBT-272: 24±4, *n*=5; prp-hTDP-43^A315T^-UeGFP+1 mg/kg/day SBT-272: 20.1±1.9, *n*=5, versus prp-hTDP-43^A315T^-UeGFP+vehicle: 59.8±7.3, *n*=4, adjusted *P*<0.0001, Fig. 8H). Interestingly, the potency of both doses to reduce astrogliosis were comparable. Administration of SBT-272 to WT-UeGFP mice did not induce astrogliosis in healthy control mice (WT-UeGFP+vehicle: 9.5±0.7, *n*=5, WT-UeGFP +5mg/kg/day SBT-272: 10.9±1.4, *n*=5, Fig. 8H).

Activated microglia accompany neuronal degeneration and exacerbate neuronal loss. Microgliosis is well documented in the motor cortex of ALS, especially within the context of TDP-43 pathology (Appel et al., 2021; Bright et al., 2021; Jara et al., 2019; Jara et al., 2017). There was also increased microgliosis in prp-hTDP-43^A315T^-UeGFP mice compared to WT healthy controls (WT-UeGFP+vehicle: 8.9±0.9, *n*=5; prp-hTDP-43^A315T^-UeGFP+vehicle: 37.1±3.7, *n*=4, adjusted *P*<0.0001). However, when prp-hTDP-43^A315T^-UeGFP mice were treated with SBT-272, the extent of microgliosis was significantly reduced at both doses (prp-hTDP-43^A315T^-UeGFP+vehicle: 37.08±3.7, *n*=4; prp-hTDP-43^A315T^-UeGFP+1mg/kg/day SBT-272: 9.1±0.6, n=5; prp-hTDP-43^A315T^-UeGFP+ 5mg/kg/day SBT-272: 13.3±2, *n*=5; adjusted *P*<0.0001, Fig. 8I, Supplementary fig. 1). Treatment of WT mice with SBT-272 did not induce gliosis, as the number of activated astrocytes or microglia remained comparable to WT healthy control levels: astrocytes (WT-UeGFP+5mg/kg/day SBT-272: 10.9±1.4, *n*=5; WT-UeGFP+vehicle: 9.5±0.7, *n*=5, *P*=0.998) and microglia (WT-UeGFP+5 mg/kg/day SBT-272: 8.9±0.7, *n*=5; WT-UeGFP+vehicle: 8.9±0.9, *n*=5, *P*=0.399). These results suggest that improving the integrity of mitochondria has a broad impact not only in improving UMN health but also reducing the extent of astrogliosis and microgliosis in the motor cortex of ALS with TDP-43 pathology.

## Discussion

ALS and other motor neuron diseases have numerous common underlying causes, such as mitochondrial defects and dysfunction. Especially problems with IMM integrity, are pronounced in ALS patients who have TDP-43 pathology (Gautam et al., 2019a) and in UMNs that become diseased due to TDP-43 pathology (Davis et al., 2018; Gao et al., 2019; Wang et al., 2013; Xu et al., 2010). Therefore, mitigating mitochondrial dysfunction is proposed to be a validated target based on clinical evidence from ALS patients with SOD1, C9orf72 mutations and with TDP-43 pathology (Filosto et al., 2011; Hervias et al., 2006; Wang et al., 2019; Wang et al., 2013), as well as recent evidence from well-characterized mouse models of ALS, which recapitulate TDP-43 pathology in patients (Gautam et al., 2019a; Wegorzewska et al., 2009).

Since SBT-272 is a small molecule in phase I clinical development with a mechanism of action that improves the integrity of IMM, we investigated whether it would have an impact on the disintegrating mitochondria of UMNs of ALS with TDP-43 pathology. We focused our attention on UMNs with TDP-43 pathology. UMNs are clinically important neurons that are in part responsible for the initiation and modulation of voluntary movement, which is impaired in ALS patients. Despite the prevailing die-back hypothesis, (Clark et al., 2016; Dadon-Nachum et al., 2011), there is now ample evidence showing UMN degeneration is an early event in ALS (Geevasinga et al., 2015; Pradhan and Bellingham, 2021; Vucic and Kiernan, 2006) and that UMN degeneration may neither be a function of spinal motor neurons loss nor depend on spinal motor neuron degeneration (Genc et al., 2022b). Improvement of UMNs health should have a direct impact on motor neuron circuitry, and potentially the health of spinal motor neurons, as well as the integrity of neuromuscular junctions (Thomsen et al., 2014), highlighting the necessity of improving UMN health, in addition to spinal motor neurons, as a treatment strategy. TDP-43 pathology that is broadly observed in ALS and ALS-frontotemporal lobar degeneration (ALS-FTLD) patients (Ling et al., 2013; Mackenzie et al., 2007; Neumann et al., 2006; Sreedharan et al., 2008), and on mitochondrial defects, which are one of the most common underlying causes of neuronal vulnerability in ALS (Filosto et al., 2011; Hervias et al., 2006; Jovicic and Gitler, 2014; Manfredi and Beal, 2000). The presence of TDP-43 proteinopathy is widely observed in a broad spectrum of ALS patients, necessitating a treatment strategy that would target mitochondrial health especially within the context of TDP-43 pathology.

UMNs with TDP-43 pathology display severe mitochondrial dysfunction both in ALS patients and in well-characterized mouse models of TDP-43 pathology (Gautam et al., 2019a). Despite species differences, UMNs in patients and CSMNs in mice have similar properties at a cellular level (Gautam et al., 2019a). This suggests the utility of investigating the cellular responses of diseased UMNs to treatment, as results would be of translational relevance to the UMNs of patients. We therefore took advantage of a well-characterized CSMN reporter line with TDP-43 pathology, in which CSMN response to compound treatment can be assayed with cellular precision.

UMNs are large layer 5 specific excitatory pyramidal neurons that extend a long apical dendrite to the top layers of the cortex, and also one of the longest axonal extensions to different segments of spinal cord targets. UMNs that are diseased due to TDP-43 pathology display massive disintegration of mitochondrial IMM in both mice and humans (Gautam et al., 2019a). The IMM is broken and disrupted, and the mitochondria are swollen in the UMNs of both species. Since the integrity of the IMM is essential for oxidative phosphorylation, a disintegrated IMM can trigger a cascade of bioenergetic crises within neurons.

SBT-272 activity is revealed only on structurally impaired mitochondria, which may occur as the result of either disease etiology (i.e., genes that impact mitochondrial structure or turnover) or because of proteinopathy leading to mitochondrial impairment or mitophagy. The potency of SBT-272 in rat cardiac cells *in vitro*, was effectively used to determine the concentration of SBT-272 (100nM) required to achieve *in vivo* pharmacodynamic effects in the renal ischemia model. The *in vivo* pharmacodynamic effect was also established in the rat brain after ET-1 induced stroke. The same concentrations were targeted to evaluate SBT-272 *in vivo* in the well-characterized TDP-43 model.

Our findings reveal that SBT-272 treatment not only improved IMM integrity but also promoted mitochondrial transport within diseased CSMN, such that mitochondria were detected in the axon and neurites in greater numbers. Since one of the cellular indications of healthy CSMN is the ability to extend a long axon and to display branching and arborization (Ozdinler and Macklis, 2006), changes in average axon length and the extent of branching/arborization are used as quantitative outcome measures to assess the health and stability of diseased CSMN. Upon SBT-272 treatment, there was an overall shift in the percentage of CSMN with longer axons and the average length of axon also significantly improved. Likewise, SBT-272 treatment also enhanced branching and arborization of diseased CSMN. Importantly, SBT-272 was efficacious and restored the overall health of impaired CSMN, so much so that it was more effective compared to edaravone and AMX0035, two drugs advanced for ALS treatment. Edaravone, a standard of care, is reported to act as a free radical scavenger (Ikeda and Iwasaki, 2015), whereas AMX0035, still in development, is the combination of two previously FDA-approved drugs that is believed to mitigate mitochondria and endoplasmic reticulum stress (Paganoni et al., 2022b). We provided evidence informing the construct validity of the model for mitochondrial dysfunction in ALS, and the results highlight the sensitivity of cultured motor neuron models to test diverse mechanisms to reduce cellular stress in ALS. The comparative pharmacology strengthens the model. While the predictive validity of the model remains to be tested in the clinic, our results suggest that SBT-272 may have therapeutically additive outcome on UMNs.

In line with cellular improvements detected *in vitro*, *in vivo* administration of SBT-272 also had a significant impact not only in improving the health of diseased UMN but also on reducing the ongoing astrogliosis and microgliosis that occur in the motor cortex of ALS with TDP-43 pathology. Recent evidence suggests that mitochondrial defects are the driving force for the initiation of innate immunity (Breda et al., 2019; Pizzuto and Pelegrin, 2020). Since mitochondrial dysfunction is known to trigger various immune reactions (Riley and Tait, 2020; West and Shadel, 2017), a link between mitochondrial health and neuroimmune reaction is suggested (Billingham and Chandel, 2019; Steinert et al., 2021; Weinberg et al., 2019). Here, we found that improvement of the health of mitochondria by chronic SBT-272 treatment resulted in profound improvements both in microgliosis and astrogliosis. These findings suggest that improvement of the health of mitochondria is upstream and that treatment strategies that focus on mitochondrial health would also have implications to reduce astrogliosis and microgliosis, two important contributors to neuronal degeneration.

## Conclusions

We find that SBT-272 treatment has a prominent impact on UMNs that are diseased due to TDP-43 pathology, especially at the site of mitochondria. SBT-272 helped regain IMM integrity and stability, enhanced mitochondrial membrane potential, and resulted in improved mitochondrial transport along the axon, which led to overall improvement of UMN health. More importantly, we also find that improving mitochondrial health leads to reduced astrogliosis and microgliosis in the motor cortex upon daily administration of SBT-272 at two different concentrations for 2 months. Since UMNs are an important component of the motor neuron circuitry that degenerates in ALS, and because TDP-43 pathology is one of the most common causes of ALS, our results highlight the potential impact of SBT-272 treatment especially within the context of mitochondrial dysfunction and TDP-43 pathology in ALS.

## Abbreviations

ADP: Adenosine Diphosphate
ALS: Amyotrophic lateral sclerosis
ANOVA: Analysis of variance
ATP: Adenosine Triphosphate
BSA: Bovine serum albumin
CHCHD3: Coiled-Coil-Helix-Coiled-Coil-Helix Domain-Containing Protein 3
CL: Cardiolipin
CLEM: Correlative light electron microscopy
CSMN: Corticospinal motor neurons
CTIP2: COUP-TF-interacting protein 2
DIV: Days in-vitro
eGFP: Enhanced green fluorescent protein
EM: Electron microscopy
ET-1: Endothelial toxin-1
FTLD: Frontotemporal lobar degeneration
GI: Gastrointestinal
IHC: Immunohistochemistry
IMM: Inner mitochondrial membrane
mSOD1: mutated superoxide dismutase 1
OMM: Outer mitochondrial membrane
OXPHOS: Oxidative phosphorylation
PFA: Paraformaldehyde
P3: Postnatal day 3
ROS: Reactive oxygen species
ROX: Residual oxygen consumption
SFM: Serum free medium
TMPD: N,N,N’,N’-Tetramethyl-p-phenylenediamine dihydrochloride
TDP-43: TARDNA binding protein 43
UCHL1: Ubiquitin C-Terminal Hydrolase L1
UMN: Upper motor neuron
VCP: Valosin-containing protein

## Author contributions

Conceptualization: M.R., D.K., M.G., P.H.O.

Methodology: B.G., M.G., P.H.O.

Investigation: B.G., M.G., N.K., A.G., B.H., P.H.O.

Funding acquisition: P.H.O.

Project administration: P.H.O.

Supervision: P.H.O.

Writing – original draft: B.G., M.G., H.Z., P.H.O.

## Acknowledgments

We would like to thank Northwestern University Center for Advanced Microscopy generously supported by NCI CCSG P30 CA060553 awarded to the Robert H Lurie Comprehensive Cancer for confocal and TEM imaging. Edijs Vavers for surgical procedures related to rat stroke model. Oge Gozutok for assistance with SBT-272 administration. The acute kidney ischemia-reperfusion study in rats was conducted at IPS Therapeutique Inc., Sherbrooke, Quebec, Canada.

## Data and materials availability

All data, code, and materials used in the analysis will be available in some form to any researcher for purposes of reproducing or extending the analysis with proper materials transfer agreements. All data is made available in the main text and the supplementary materials.

## Supplementary Materials

**Supplementary table S1:** Pharmacokinetic properties of SBT-272.

**Supplementary table S2:** Statistics for EM analysis shown in Fig. 3.

**Supplementary table S3:** Mean length of the longest neurite for each treatment.

**Supplementary table S4:** Statistics for Sholl analysis shown in Fig. 7.

---

**Supplementary fig. S1:**
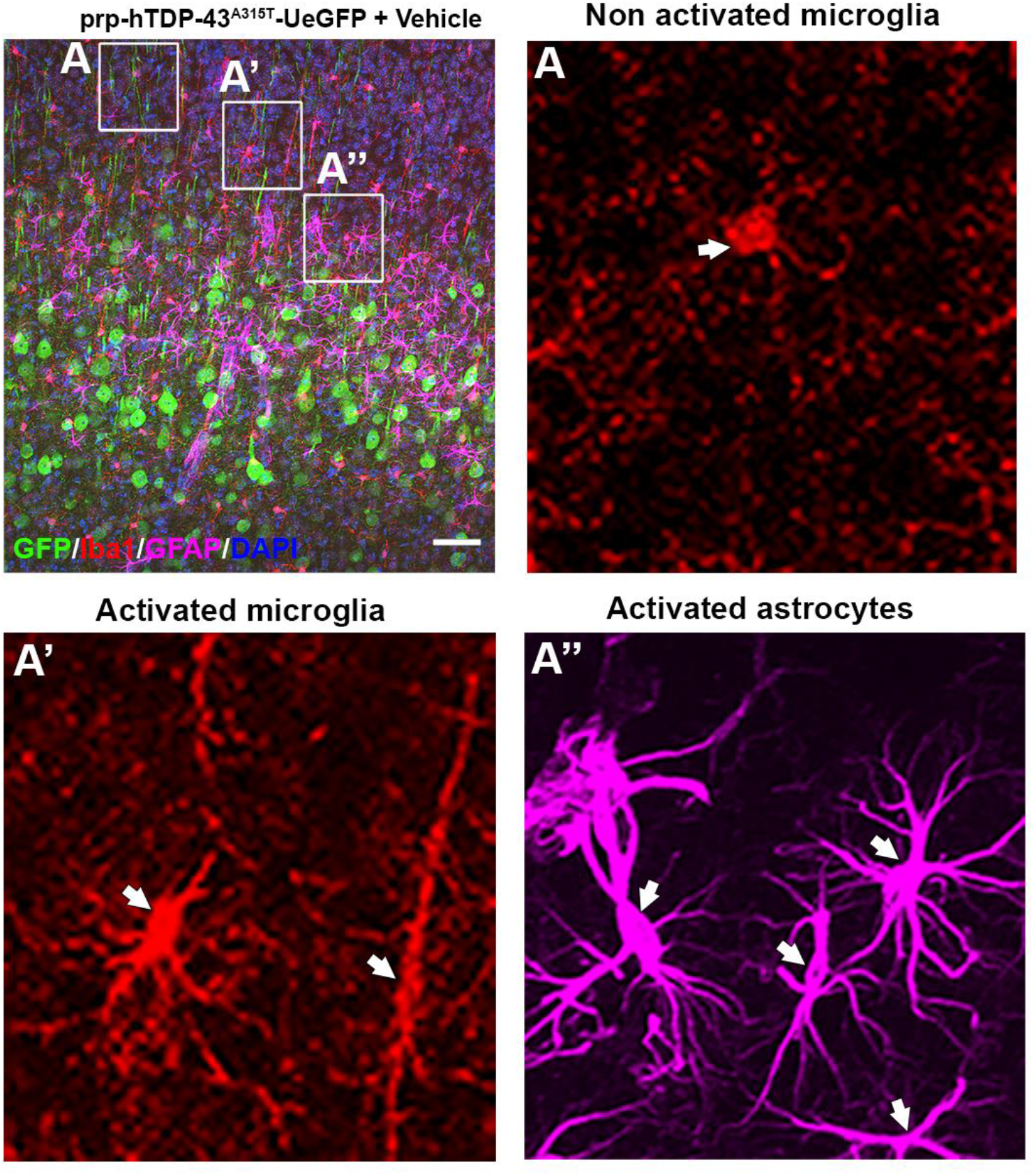
Activated astrocytes and microglia are identified based on their morphology and the markers they express. Scale bar: 50μm. Arrows in (A), (A’) and (A”) point to non-activated microglia, activated microglia, and activated astrocytes respectively.

