## Supplementary table 1 and 3 for "SBT-272 improves TDP-43 pathology in the ALS motor cortex by modulating mitochondrial integrity, motility, and function"

**Supplementary Materials:**

**Supplementary Tables:**

|  | Plasma | Brain |
| --- | --- | --- |
| C_max_ | 5017ng/ml | 201ng/g |
| t_1/2_ | 3.34 | ND |
| AUC_0–24h_ | 16283ng/ml•h | 3732ng/g•h |
| Tissue/plasma AUC ratio | _ | 22.9% |

**Supplementary table S1. Pharmacokinetic properties of SBT-272.** AUC, area under the concentration-time curve; AUC_0–24h_, AUC from time zero to 24 hours; C_max_, maximum concentration; ND, not determined; t_1/2_, half-life.

| Treatment | Mean (±SEM)  axon length (µm) | Mice (*n*) | CSMNs (*n*) |
| --- | --- | --- | --- |
| SFM | 154.68 (5.99) | 6 | 77 |
| SBT-272 (10nM) | 360.09 (20.58) | 3 | 122 |
| SBT-272 (100nM) | 418.88 (25.88) | 6 | 108 |
| SBT-272 (1µM) | 291.63 (23.46) | 3 | 45 |
| Edaravone (10nM) | 196.28 (8.63) | 3 | 58 |
| Edaravone (100nM) | 244.83 (4.70) | 3 | 48 |
| Edaravone (1µM) | 279.21 (13.57) | 6 | 191 |
| AMX0035 (1mM sodium phenylbutyrate + 100µM taurursodiol) | 271.66 (18.66) | 3 | 63 |

**Supplementary table S3.** Mean length of the longest neurite for each treatment.
